# Comparing colours using visual models

**DOI:** 10.1101/175992

**Authors:** Rafael Maia, Thomas E. White

## Abstract

**Lay Summary:** An outstanding challenge for the study of colour traits is how best to use “colour spaces” to represent their visual perception, particularly when asking questions of colour-difference (e.g. the (dis)similarity of males and females, mimics and models, or sister species, to a given viewer). We use simulations to show that existing methods fail to statistically and biologically estimate the separation of groups in colour space, and we suggest a flexible, robust, alternative that avoids those pitfalls.

**Abstract:** Colour in nature presents a striking dimension of variation, though understanding its function and evolution largely depends on our ability to capture the perspective of relevant viewers. This goal has been radically advanced by the development and widespread adoption of colour spaces, which allow for the viewer-subjective estimation of colour appearance. Most studies of colour in camouflage, aposematism, sexual selection, and other signalling contexts draw on these models, with the shared analytical objective of estimating how similar (or dissimilar) colour samples are to a given viewer. We summarise popular approaches for estimating the separation of samples in colour space, and use a simulation-based approach to test their efficacy with common data structures. We show that these methods largely fail to estimate the separation of colour samples by neglecting (i) the statistical distribution and within-group variation of the data, and/or (ii) the discriminability of groups relative to the observer’s visual capabilities. Instead, we formalize the two questions that must be answered to establish both the statistical presence and theoretical magnitude of colour differences, and propose a two-step, permutation-based approach that achieves this goal. Unlike previous methods, our suggested approach accounts for the multidimensional nature of visual model data, and is robust against common colour-data features such as heterogeneity and outliers. We demonstrate the pitfalls of current methods and the flexibility of our suggested framework using an example from the literature, with recommendations for future inquiry.

## Introduction

The study of colour in nature has driven fundamental advances in ecology and evolutionary biology (Cuthill *et al*., 2017). Colour is a subjective experience, however, so substantial effort has been dedicated to measuring and representing colours “objectively” (Garcia *et al*., 2014; Johnsen, 2016) through visual models that consider the perspective of ecologically relevant viewers (Kemp *et al*., 2015; Renoult *et al*., 2017). These models have significantly advanced the study of colour traits by allowing researchers to account for the factors influencing the generation and reception of visual information, such as the structure of signals and viewing backgrounds, the properties of veiling and incident light, and the attributes of visual systems (Chittka, 1992; Endler & Mielke, 2005; Kelber *et al*., 2003; Vorobyev & Osorio, 1998).

Several forms of visual models are currently used, which vary in their assumptions about the nature of visual processing (Chittka, 1992; Endler & Mielke, 2005; Vorobyev & Osorio, 1998). These models function by delimiting a colour space informed by the number and sensitivity of photoreceptors in an animal’s retina (Renoult *et al*., 2017). Individual colours are then represented in this space as points, with their location determined by the differential stimulation of the viewers’ receptors.

This colour space representation is convenient for several reasons. It offers an intuitive way of analysing phenotypes that we cannot measure directly: we can estimate how animals with different visual systems “see” different colours by representing them in a Cartesian coordinate system, producing a receiver-dependent morphospace (Kelber *et al*., 2003; Renoult *et al*., 2017). Further, it allows estimating how similar or dissimilar colours are *to a given observer*, by measuring the distance between colour points in its colour space (Endler & Mielke, 2005; Vorobyev *et al*., 1998; Vorobyev & Osorio, 1998). Crucially, we can test and refine these models using psychophysical data (e.g. Dyer & Neumeyer, 2005; Garcia *et al*., 2017;

Maier, 1992; Vorobyev *et al*., 2001), to estimate the magnitude of colour-differences and ultimately predict whether an observer could effectively discriminate pairs of colours (Chittka, 1992; Vorobyev & Osorio, 1998). This final point is critical to many tests of ecological and evolutionary hypotheses, such as the efficacy of camouflage (Pessoa *et al*., 2014; Troscianko *et al*., 2016), the presence of polymorphism or dichromatism (Schultz & Fincke, 2013; Whiting *et al*., 2015), the accuracy of mimicry (O'Hanlon *et al*., 2014; White *et al*., 2017), the extent of signal variability among populations or species (Delhey & Peters, 2008; Rheindt *et al*., 2014), or the effect of experimental manipulations (Barry *et al*., 2015; White & Kemp, 2017). At the heart of these inquiries lies the same question: *how different are these colours to the animal viewing them?*.

### Challenges in estimating the discriminability of colour samples

The receptor noise-limited model of Vorobyev & Osorio (1998) has proven particularly useful for addressing questions of discriminability and colour-difference. The model is focused on receptor-level processes, and assumes that chromatic and achromatic channels operate independently (which does not necessarily hold beyond the receptor level in some species, such as humans; Nathans, 1999), that colour is coded by *n —* 1 unspecified opponent mechanisms (where *n* is the number of receptor channels), and that the limits to colour discrimination are set by noise arising in receptors (Vorobyev *et al*., 1998; Vorobyev & Osorio, 1998). This noise is dependent on the receptor type and abundance on the retina which, along with Weber’s law (*k* = Δ*I/I*) more generally, ultimately establishes the unit of Just Noticeable Differences (JND Vorobyev *et al*., 2001). Distances calculated in this manner correspond to the Mahalanobis Distance *_DM_*, and represent distances between points standardized by the Weber fraction; i.e. 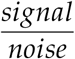 (Clark *et al*., 2017). It follows that values lower than 1 JND 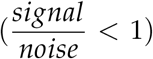 are predicted to be indistinguishable, while values greatly above this threshold are likely distinct. This provides a useful standard for estimating the similarity of groups of points in colour space: the greater the distance between colours, the less alike they are. If differences are, on average, above an established threshold, then we can consider the groups different: sexes dichromatic, mimetism imperfect, crypsis ineffective. This offers a clear link between variation and classification within a sensory framework, and has been widely used for this purpose (Barry *et al*., 2015; Delhey & Peters, 2008; O’Hanlon *et al*., 2014; Schultz & Fincke, 2013; White *et al*., 2017; White & Kemp, 2017).

To adequately compare samples of colours, however, it is necessary to determine if the average distance between them is both statistically and biologically meaningful (i.e. above-threshold; Endler & Mielke, 2005). Commonly, an “average colour” for each group is derived by taking a mean reflectance spectrum or by averaging their position in colour space. In either case, the colour distance between groups is then calculated from these mean quantum catches per-receptor per-group — their centroids in multivariate space (Fig. 1, bold arrow). However, the centroid obtained from arithmetic means of receptor coordinates is not an appropriate measure of location for this purpose, since colour distances are perceived in a ratio scale (Cardoso & Gomes, 2015). Instead, the geometric mean must be used. Further, since the result is a single value representing the multivariate distance between group means, there is no associated measure of uncertainty or precision that would allow for the statistical testing of differences between samples (e.g. Avilés *et al*., 2011; Burns & Shultz, 2012; Maia *et al*., 2016).

An alternative approach calculates the pairwise distances between all points in group A and group B, then averages these distances to obtain a mean distance between groups (Fig. 1, thin arrows; e.g. Barry *et al*., 2015; Dearborn *et al*., 2012). In cluster analyses, this is called the “average linkage” between groups (Hair *et al*., 1998). This is an appealing method, providing measures of variation among distances, and thus a t-test or equivalent can be used to test if differences are greater than a given threshold. The average linkage, however, is also inadequate because it conflates within- and among-group variation. This is because Euclidean distances (and by extension JND’s) are *translation-invariant:* they ignore the position of points in colour space and the direction of the distance vectors, reflecting only the magnitude of differences between two points. Therefore, the average linkage reduces to a measure of spread, and will scale with both within- and between-group distances (Fig. 1, insert).

As these issues show, hypotheses of discriminability and colour-difference have primarily focused on testing whether the difference between samples is above a theoretical threshold. However, the convenience of such thresholds belies fact that simply comparing means between groups is not sufficient to infer, statistically, whether samples are different. To answer if two groups are different, one must compare the variation between- and within-groups. This is particularly problematic in the case of colours that function as signals in social interactions (e.g. Kemp & Rutowski, 2011). For a trait to function in this context, the observer must be able to tell signals of ‘low’ and ‘high’ quality apart. This means that, by definition, *most pairs of individuals should be readily distinguishable*. The trait must be highly variable and colour distances should be above the threshold of discrimination (Delhey *et al*., 2017), otherwise no information can be extracted by an observer comparing phenotypes.

Consider a hypothetical species that uses colour in mate choice, but is not sexually dichromatic (Fig. 1). In this species colour is highly variable and, on average, pairs of individuals are discriminable, but there is no *consistent* male-female difference. Therefore, if a researcher sampled this species and calculated the average distance between all pairs of individuals, regardless of sex, these should be largely greater than 1 JND. However, if they took separate samples of males and females, then all pairwise distances (the average linkage) between sexes will be also greater than 1 JND, despite them being sampled from the same (statistical) population.

**Figure 1:**
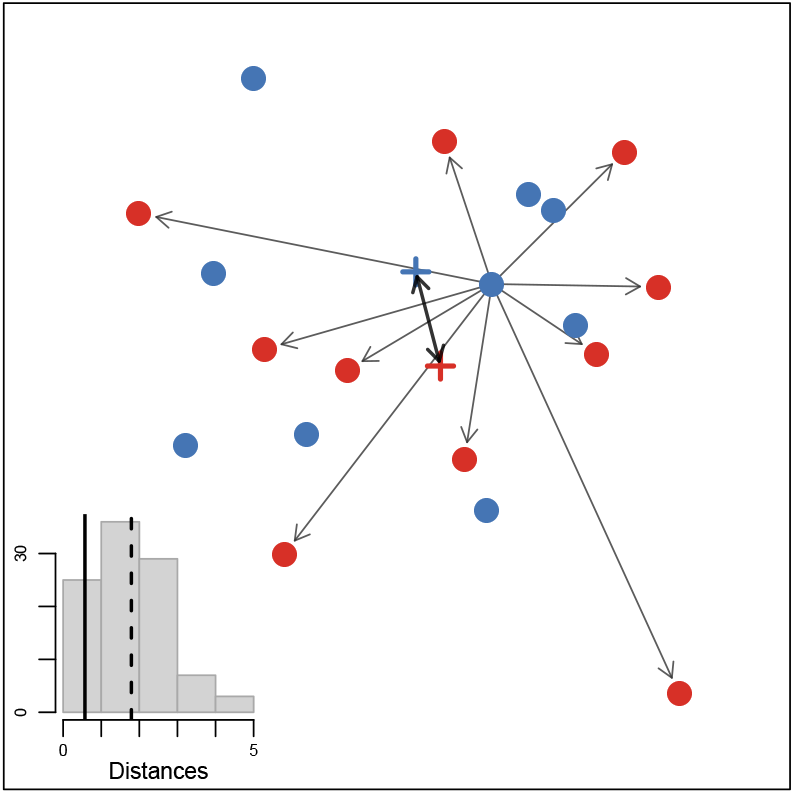
The link distance (i.e. average pairwise distance between groups) conflates within- and among-group variation. Here, two samples were drawn from the same simulated distribution. Thin arrows represent distances between a random point in the first sample (blue) and all points from the second sample (red), all of which are greater than the distance between the geometric means of the two samples (“x”, bold arrows). Inset shows the histogram of pairwise distances among groups, and how their average (dashed line) is greater than the mean distance (bold line).

### The limitations of current methods for comparing colour space distributions

Several methods have been proposed to avoid the aforementioned issues by accounting for the relative distributions of samples in colour space. Eaton (2005), for example, noted that within-group variation influenced the conclusions on the extent of avian dichromatism, and thus tested for intersexual differences in photon catches separately for each receptor. However, this ignores the multivariate nature of visual model data by failing to account for multiple comparisons and correlations among receptor catches (which are critical, since any *n*-receptor visual system can be represented in *n —* 1 dimensions; Kelber *et al*., 2003).

An alternative, multivariate metric suggested by Stoddard & Prum (2008) is the volume overlap. In this approach, the volume occupied by a sample of colours is estimated from its enveloping convex hull, and separation between samples is inferred from their overlap. Stoddard & Stevens (2011) used this metric to show that a greater overlap in colour volume between cuckoo and host eggs is associated with lower rejection of parasitic eggs. This approach is appealing because it considers the distribution of colour points in multivariate space, though there are limits to its interpretation: (i) there is a lower bound to group separation (i.e. if samples do not overlap, there is no distinction between cases where samples are near or far apart) and (ii) it is unclear how variation in volume overlap should be interpreted biologically (e.g. how biologically relevant is the difference between 20% or 40% overlap?). It is also particularly sensitive to outliers, because the volume defined by a convex hull does not lend itself to a probabilistic interpretation, leading to the often unacknowledged assumption that the sampled data reflects the true boundaries of the population (however, “loose wrap” hypervolumetric methods exist; to our knowledge, these have not been applied to colour studies; Blonder *et al*., 2017). Finally, in its original implementation this method does not consider receptor noise or discrimination thresholds (but incorporating this is straightforward; see below).

The most robust attempt at comparing distributions of colours was proposed by Endler & Mielke (2005), who devised a non-parametric rank distance-based approach based on the least sum of Euclidean distances, compared through multiresponse permutation procedures (LSED-MRPP). This multivariate approach is powerful because it calculates an effect size based on the relationship of between- and within-group distances. However, this single statistic captures differences between samples not only in their means, but also in their dispersion and correlation structure (i.e. shape; Endler & Mielke, 2005). Like other distance-based methods, it is sensitive to confounding heterogeneity among samples when testing for differences in *location* (Anderson & Walsh, 2013; Warton *et al*., 2012). Despite its considerable strengths, this method has seen little adoption over the last decade, largely due to limitations in implementation and accessibility.

The shortcomings of these methods reflect the fundamental fact that the question of discriminability actually represents a test of two hypotheses that are seldom formally distinguished: (i) that the focal samples are statistically distinct, and (ii) that the magnitude of their difference is greater than a psychophysical threshold of detection. Most approaches will test one, but not both, of these hypotheses through their respective nulls, and often with no estimate of variation or uncertainty in estimates. We explore these issues using a simulation-based approach by testing the efficacy of popular methods in detecting the separation of groups in colour space. We then propose a flexible solution that avoids these problems, demonstrating its utility using an example from the literature.

## Methods

### Simulation procedures

To compare methods for detecting group separation in colour space, we simulated data analogous to that obtained from applying an avian visual model to spectral reflectance data. Birds are tetrachromatic (Hart, 2001), and colours will thus be represented by the quantum catches of its four photoreceptors (though the procedure followed here can be applied to visual systems with any number of receptors). For each replicate, we simulated two samples defined by four variables (*USML* photoreceptors) taken from log-normal distributions (since quantum catches are non-negative and noise-corrected distances follow a ratio scale, as defined by the Weber fraction, described above). We generated samples following two different scenarios: first, we simulated varying degrees of separation (i.e. effect sizes) to evaluate the power and Type I error rates of the approaches tested. Second, we simulated threshold conditions to evaluate the performance of different approaches in correctly classifying whether samples are above-threshold.

For the first set of simulations (power and error-rates) we simulated the quantal catch of each photoreceptor *i* for the first sample (group A) by drawing from a log-normal distribution with mean *μ¡A* seeded from a uniform distribution 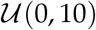, and standard deviation proportional to the mean: 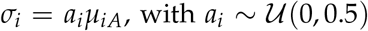 (note that, for these simulations, *μ* and *σ* refer to the mean and standard deviation of the random variable itself, not in log scale). To generate two samples with varying degrees of separation proportional to the within-group variance, we used a multivariate effect size *S* obtained by calculating a constant 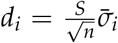, where *n* is the number of photoreceptors (in this case, 4) and 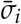 is the standard deviation of the sample. We then drew a second sample (group B) defined by *μ_iB_* = *μ_iA_* + *d_i_* and *σ_i_*. Thus, our simulations effectively produced two samples with Mahalanobis Distance *D_m_* ~ *S* (calculated as the distance between centroids of the two groups weighted by their pooled variance-covariance matrix). We simulated data for *S* = {0,0.1,0.25,0.5,0.75,1.0,1.5,2,2.5,3.0} (Fig. 2), replicated 200 times for sample sizes *N =* {10,20,50,100} each.

**Figure 2:**
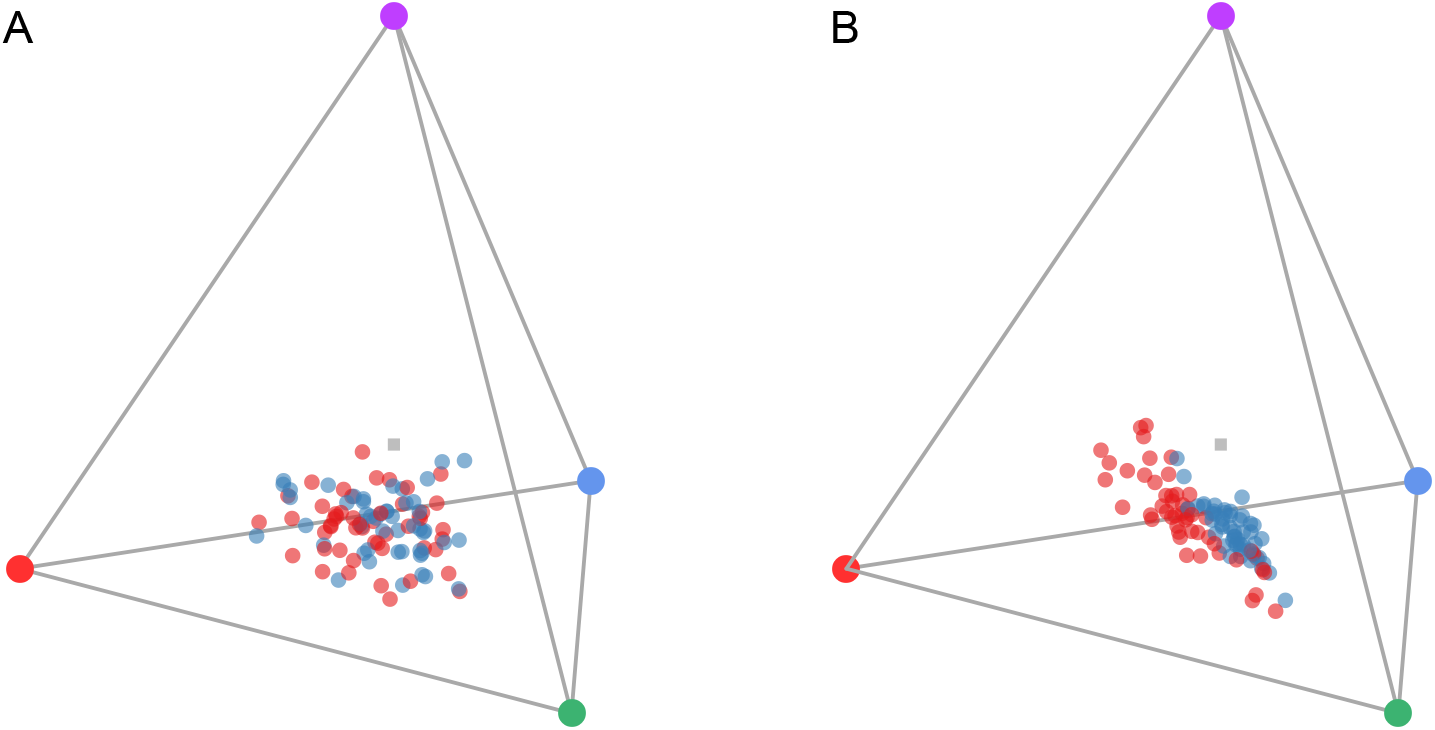
Example simulated data for the two groups (red, blue) in a tetrahedral colourspace. Shown here are data with sample size *N* = 50 and effect size (A) *S* = 0 and (B) *S* = 3.

For the second set of simulations (threshold conditions across a range of within-sample variation), we followed a similar procedure. Group A was sampled from a log-normal distribution with 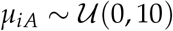, while σ was taken from an exponential distribution *σ_i_* ~ *Exp*(*λ* = 1). To obtain a second sample, group B, that was separated from group A with an average approximate distance of ~ 1 JND given a Weber fraction of 0.1 for the long-wavelength photoreceptor (Vorobyev *et al*., 1998), we drew from log-normal distributions with *μ_iB_* = *d_i_μ_iA_*, where 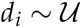(0.88,1.12), resulting in an average distance between geometric means (hereafter, “mean distance”) of 1.11 (95% quantiles: 0.35 — 2.77 JND) and within-group average pairwise distance of 4.46 (95% quantiles: 1.03 — 11.10 JND) after 1000 replicates.

After the two groups were simulated, we used the R package pavo (Maia *et al*., 2013) to calculate colour distances using relative receptor densities of {*U, S, M, L*} = {1,2,2,4} and Weber fraction for *L* = 0.1. We calculated the within-group average pairwise distance, as well as the distance between sample geometric means.

We then used four procedures to test for differences between groups. First, we used a distance-based PERMANOVA (hereafter “distance PERMANOVA”) using the adonis function from the R package vegan (Oksanen *et al*., 2007). This non-parametric approach uses distances to calculate a pseudo-F statistic, simulating a null distribution by randomizing distances between observations (Anderson, 2005). We recorded if the analysis was significant (α = 0.05) using 999 permutations for the null, as well as the *R*^2^ as an effect size estimate. Second, we obtained *XYZ* Cartesian coordinates based on “perceptually-scaled” (i.e. noise-corrected) distances (Pike, 2012; functionally and mathematically equivalent to the receptor-noise limited space of de Ibarra *et al*., 2001) and applied a MANOVA test on these coordinates (hereafter “Cartesian MANOVA”). For simplicity, we used a sum of squares and cross-products matrix approach and calculated Pillai’s trace and its associated P-value (but see discussion for extensions of this approach). Third, we calculated the volume overlap between the two samples (relative to their combined volumes) in a tetrahedral colour space defined by the receptors’ relative quantum catches (thus disregarding receptor noise; Stoddard & Prum, 2008). Finally, we calculated the volume overlap for the *XYZ* Cartesian coordinates based on noise-corrected distances, generating a colour volume overlap that accounts for receptor noise.

## Simulation results

### Power and error rates

Both the distance PERMANOVA and the Cartesian MANOVA showed appropriate Type-I error rates, with about 5% of our simulations producing significant results when *S* = 0, even for small sample sizes (Fig. 3). As expected, the power to detect small effects steadily increased as a function of sample size, with the distance PERMANOVA being overall more conservative than the Cartesian MANOVA (Fig. 3,4).

**Figure 3:**
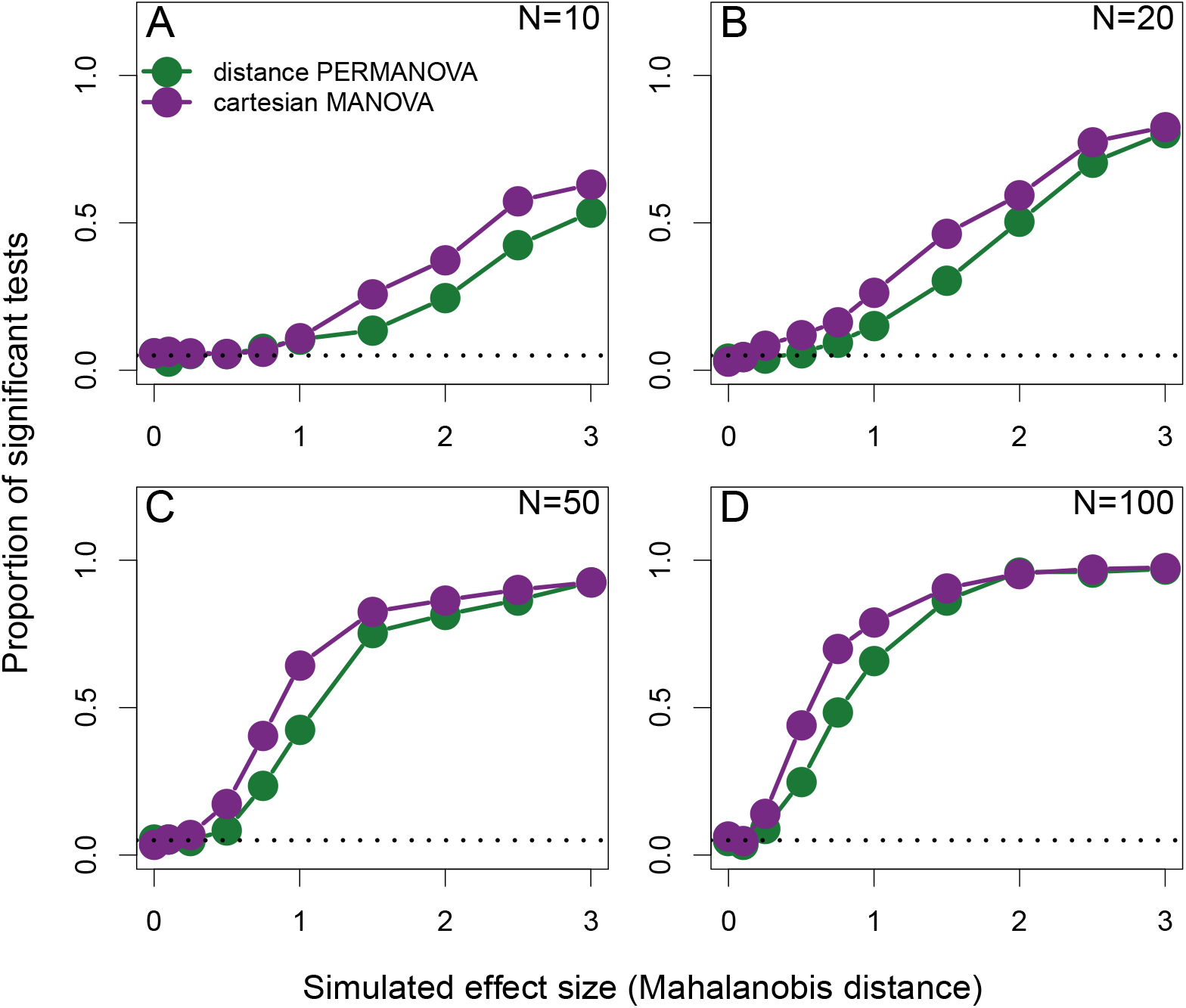
Power and Type I error rate of the distance PERMANOVA (green) and Cartesian MANOVA (purple). Panels show the proportion of simulations yielding significant results for each approach under different sample and effect sizes.

The two approaches showed some disagreement, with between 10 — 15% of the simulations significant only in one of the two tests (Fig. 4). This disagreement was not random, with the Cartesian MANOVA being more likely to be significant when the distance PERMANOVA was not than vice-versa (Fig. 4a), at an approximately constant rate across sample sizes, and disagreemennt being concentrated at smaller effect sizes with increasing sample sizes (Fig. 4b).

**Figure 4:**
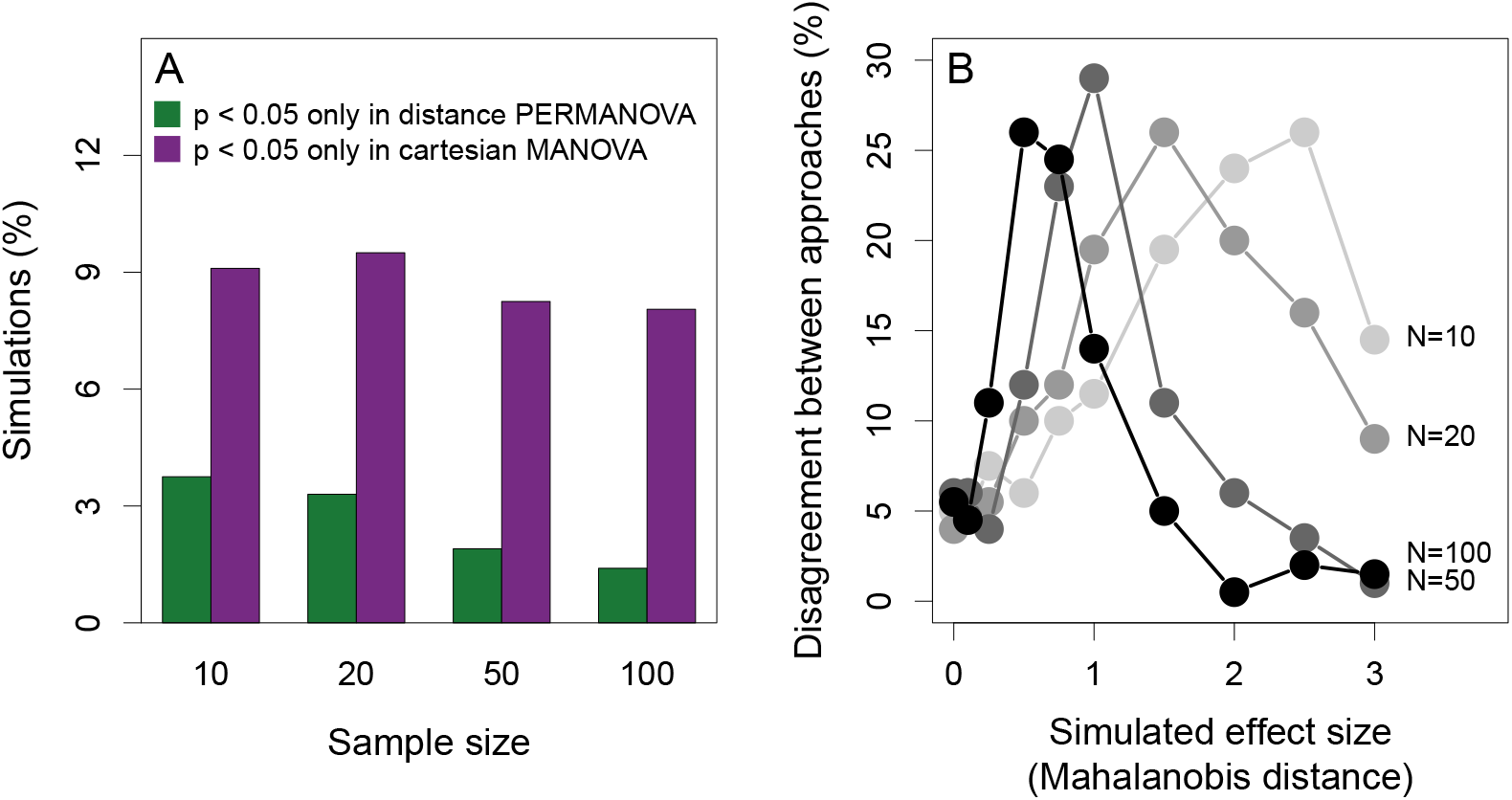
The disagreement between multivariate statistical approaches when testing for separation between samples in colour space in relation to sample size (A) and effect size (B).

Focusing on *N* = 50 simulations, our results show that mean distance was positively associated with the effect size, and the threshold of significance using the distance PERMANOVA fell approximately at the 1 *JND* mark (Fig. 5A; equivalent results are observed with the Cartesian MANOVA, not shown). Still, even around that threshold, significance is variable, showing that large within-group variation can lead to non-significant differences between groups despite among-group distances being above the theoretical perceptual threshold. Volume overlap also showed a (negative) association with effect size, but no specific threshold for significance is observed (e.g. both significant and non-significant results are observed for 20 — 60% overlap; Figure 5B).

**Figure 5:**
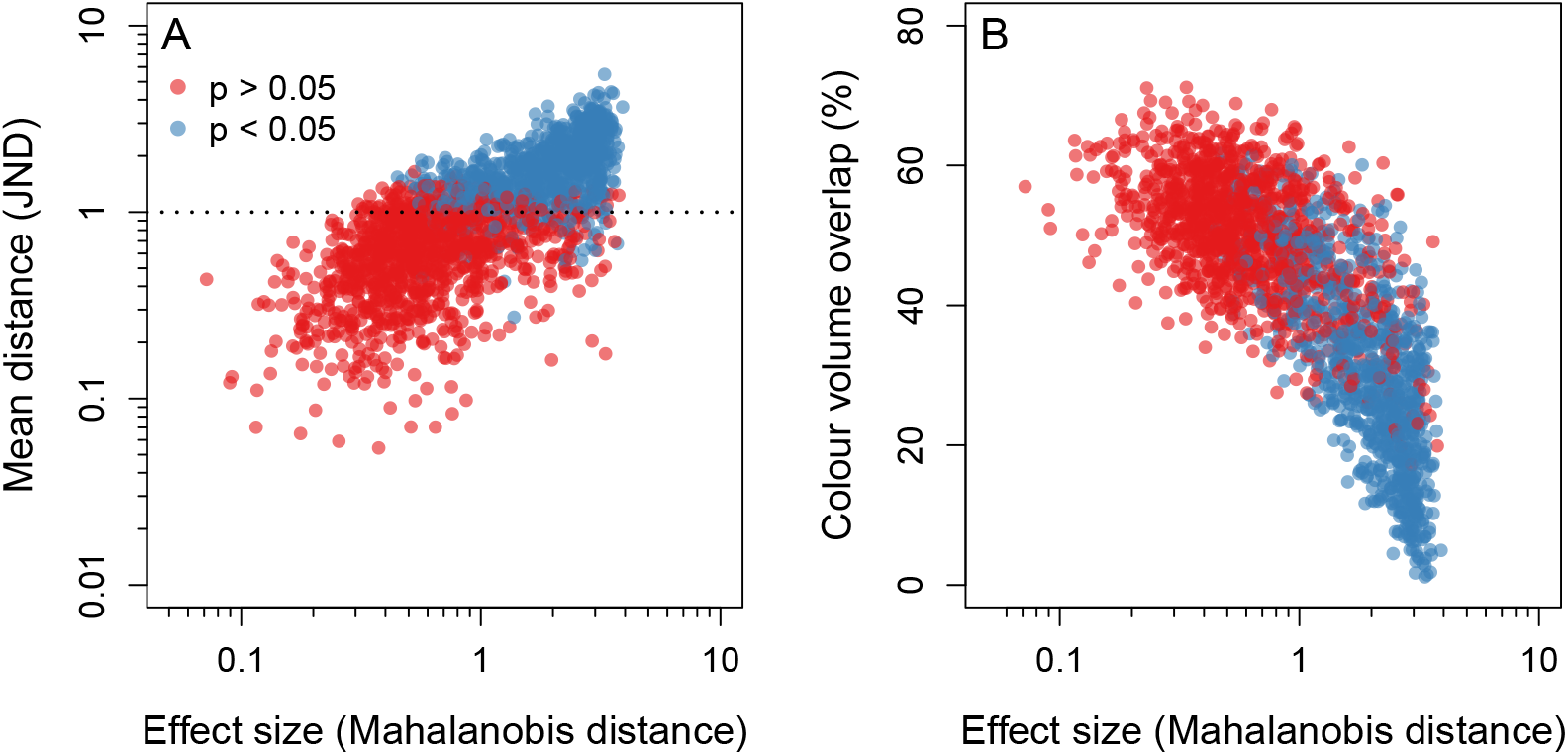
The association between effect size and (A) mean distance and (B) colour volume overlap. Significant distance PERMANOVA results are in blue, whereas non-significant results are in red. Dotted line indicates the threshold of 1 JND.

### Threshold scenarios

Since results from the distance PERMANOVA and the Cartesian MANOVA were comparable, we focus on the former due to the convenience of the *R*^2^ statistic describing among-group separation (but see Discussion for comments on the use of these approaches). Simulations produced a wide range of outcomes, with nonsignificant and significant tests both above and below the theoretical threshold of 1 JND (Fig. 6). In contrast with the power simulations above (Fig. 5), the significance threshold did not match the theoretical perceptual threshold. As in the hypothetical example from the introduction, 20.2% of the simulated cases were statistically indistinguishable despite having mean above-threshold distances (Fig. 6, dark red). Likewise, 15.1% of the simulations produced samples that were statistically different, but where this difference was below threshold and was therefore likely undetectable to its observer (Fig. 6, dark blue points). These results highlight the importance of considering both statistical separation and theoretical perceptual thresholds when testing the hypothesis that samples are discriminable.

**Figure 6:**
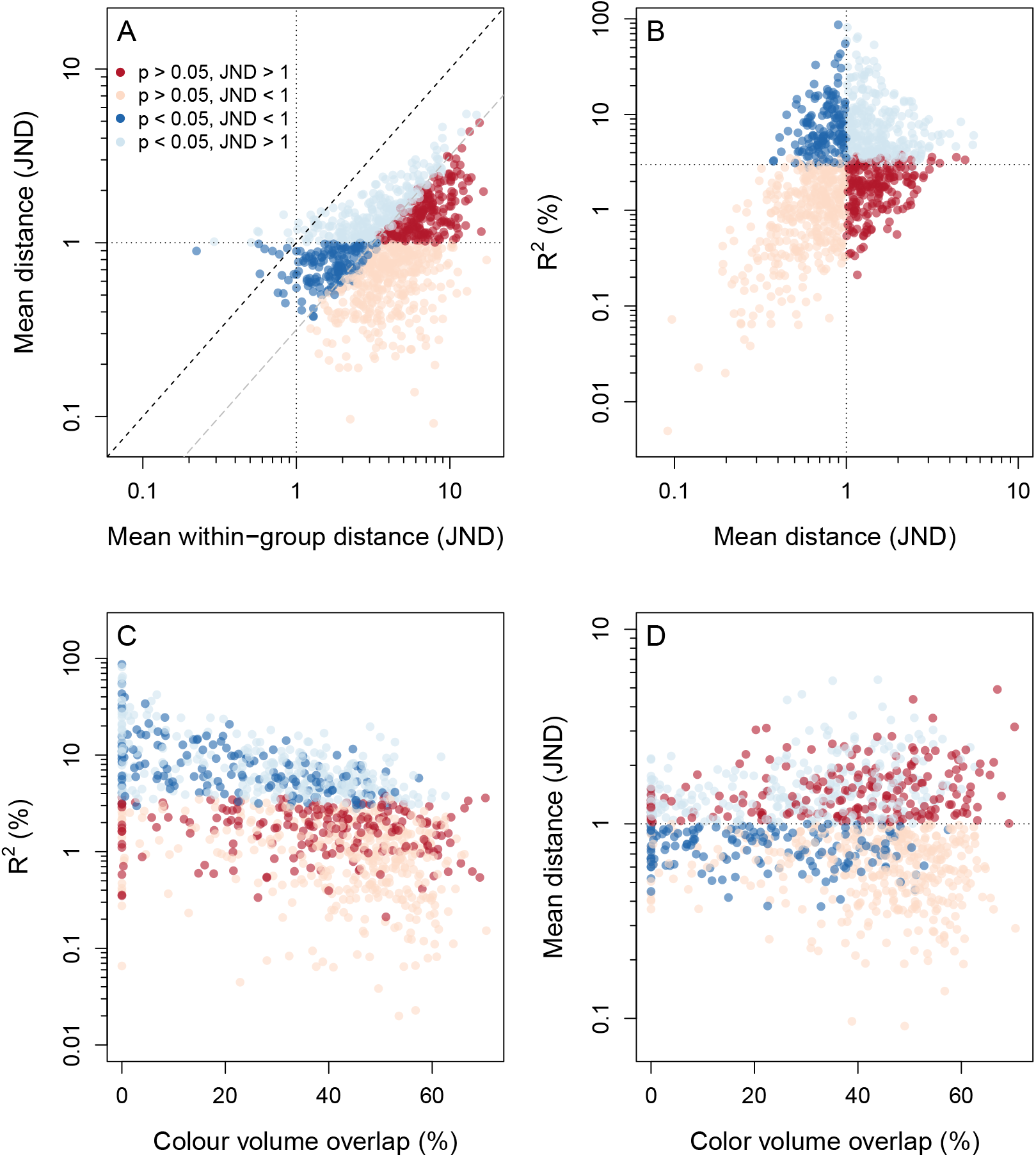
Results from threshold simulation. Red and blue denote non-significant and significant PERMANOVA tests, respectively, and light colours denote when that approach would yield the same inference as comparing mean distances to a threshold of 1JND. Thus, dark blue points indicate a significant statistical test that does not reach the threshold of discriminability of 1 JND, whereas dark red points indicate a non-significant statistical test that nonetheless has a mean distance greater than 1 JND.

Figure 6A shows that, intuitively, tests were significant when within-group differences were small relative to among-group differences. However, nearly all simulations — including most significant results — fell below the 1:1 line when using the average link distance (i.e. the average pairwise distance) to describe intragroup variation. Significant results are obtained when the mean difference is up to 0.5 JND smaller than the within-group average link distance (Fig. 6A, grey line intercept). Similarly, we can see that significant results can be obtained for fairly low levels of among-group separation, with *R*^2^ as small as 3 or 4% (Fig. 6B, horizontal line at 3%).

Though there is a negative association between *R*^2^ and volume overlap (Fig. 6C), results show low overall consistency between approaches: for any given value of volume overlap, all possible outcomes of significance/threshold occur — even when the overlap between samples is zero (Fig. 6C). In other words, even complete separation in colour volumes can result in non-significant, below-threshold cases, since samples can be contiguous without overlapping in noise-corrected colour space. Likewise, samples can have high overlap but be entirely distinguishable statistically and perceptually. Further, there is no association between volume overlap and mean distance between groups (Fig. 6D). These results were unaltered by considering receptor noise in the volume overlap calculation, since these are still strongly and positively correlated with their non-noise-corrected counterparts (Electronic Supplementary Material.

### A two-step approach to estimate statistical and perceptual separation

As described previously, questions of discriminability and colour-difference require testing two distinct hypotheses: if samples are (i) statistically and (ii) ‘perceptually’ distinct. We therefore propose a two-step answer to such questions, which explicitly formalizes these hypotheses. For the first question — are the samples *statistically separate* in colour space? — we show that both a PERMANOVA using noise-corrected colour distances (Anderson, 2005; Cornuault *et al*., 2015), and a MANOVA using noise-calibrated Cartesian coordinates (de Ibarra *et al*., 2001; Delhey *et al*., 2015; Pike, 2012) are well suited. Both exclude achromatic variation and properly account for the multivariate nature of the data. There is also minimal discrepancy between the two (Fig. 3,4), so the decision between them may be informed by convenience and the structure of the data at hand.

Once the separation of samples is established statistically, a second question must be answered: is this separation predicted to be *perceptually discriminable*? The statistics calculated above cannot answer this, since effect sizes account for both among- and within-group variance. We therefore suggest this be tested independently, by estimating the distance in colour space between group geometric means rather than through the average pairwise distance or volume-overlap based metrics, which fail to accurately estimate group separation (Figs. 1,6). One limitation to this statistic is the lack of any measure of uncertainty. To circumvent that, we suggest a bootstrap procedure in which new samples are produced through resampling (with replacement) of individuals of each group, from which geometric means and their distance are calculated. Repeating this procedure generates a distribution of mean distances, from which a confidence interval can be estimated. If the groups being compared are statistically different and this bootstrapped confidence interval does not include the theoretical threshold of adequate biological significance, one can conclude that the samples being compared are distinct and likely discriminable.

## Empirical example: Sexual dichromatism in the leaf-nosed lizard *Ceratophora tennentii*

Visually signalling animals often use distinct body parts for different purposes, such as social signalling to mates or warning predators (Barry *et al*., 2015; Grether *et al*., 2004; Johnstone, 1995). The nature of intraspecific variation in colour can thus inform their putative function, since selection may act differentially on signals used in different contexts. For example, traits subject to strong sexual selection in one of the sexes are often dimorphic, with one sex (typically males) expressing a conspicuous colour pattern that is reduced or absent in the other (Bell & Zamudio, 2012; Kemp & Rutowski, 2011).

Dragon lizards (Agamidae) are known for variable colouration used in both social and anti-predator contexts (Johnston *et al*., 2013; Somaweera & Somaweera, 2009). The leaf-nosed lizard *Ceratophora tennentii* has multiple discrete colour patches, with apparent sex differences between body parts (Fig. 7). Here we draw on the data of Whiting *et al*. (2015), who recorded the spectral reflectance of 29 male and 27 female *C. tennentii* from four body regions (throat, labials, mouth-roof, and tongue). We used a tetrachromatic model of agamid vision to test for dichromatism among body regions from the perspective of conspecifics.

Following standard calculations for the log-linear receptor-noise model, we used the spectral sensitivity of *Ctenophorus ornatus (λ_max_ =* 360,440,493,571 nm) as modelled according to a vitamin A1 template (Barbour *et al*., 2002; Govardovskii *et al*., 2000). We assumed a relative photoreceptor abundance of 1: 1: 3.5: 6, and a coefficient of variation of noise yielding a Weber fraction of 0.1 for the long-wavelength cone (Fleishman *et al*., 2011; Loew *et al*., 2002). We tested each body region separately using PERMANOVA. As above, we used the R package pavo for visual modelling, and the adonis function in the R package vegan for PERMANOVAs.

We found a statistical difference between male and female throats (PERMANOVA: F_1,58_ = 14.84, *P* < 0.01) and labials (*F*_1,57_ = 13.96, *P* < 0.01; Fig. 7A,B), but not for tongues (F,_1,58_ = 1.63, *P* = 0.22) or mouth-roofs (*F*_1,55_ = 0.52, *P* = 0.50; Fig. 7C,D). However, bootstraps of group separation suggest that intersexual differences in labial colour are likely imperceptible to conspecifics (Fig. 7E; though like all such predictions this requires behavioural validation). Our results therefore suggest the absence of dichromatism in all but throat colour from the lizard perspective, despite statistical significance for the labial region. These results thus do not implicate sexual selection as a strong driver of intersexual colour differences in these few body regions of *C. ornatus*.

**Figure 7:**
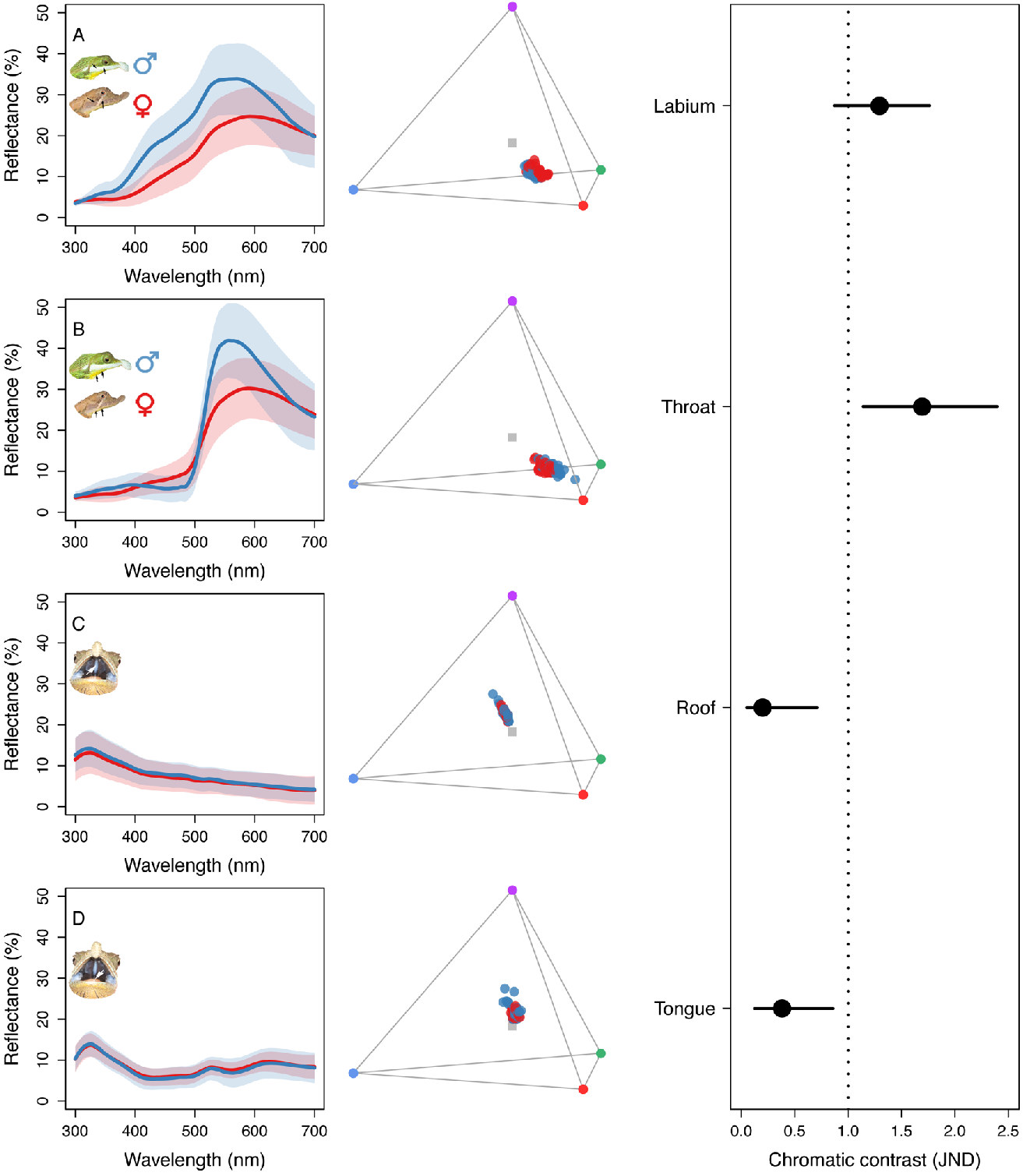
The mean (± SD) spectral reflectance of female (red) and male (black) (A) labial, (B) throat, (C) mouth-roof, and (D) tongue (left panels), and their colourspace distribution according in a tetrachromatic model of agamid vision (middle panels). Inset images indicate approximate sampling regions. The bootstrapped 95 % C.I’s for mean distances between groups in colour space (right panels). Partly reproduced, with permission, from Whiting *et al*. 2015.

## Discussion

Visual models offer a useful tool for quantifying the subjective perception of colour, which — as the ultimate canvas for colour-signal evolution — can offer valuable insight into a breadth of biological phenomena. It is therefore essential that statistical considerations of biological hypotheses take into account both natural variation in the compared samples as well as the limits to observer perception (as ultimately informed by behavioural and physiological data; Kemp *et al*., 2015). Here, we show that most methods typically fail to consider these aspects, and propose a flexible alternative that explicitly addresses both.

The use of models that do not explicitly consider discriminability, such as the volume-overlap and segment-based analyses, is often justified on the basis of simplifying and relaxing assumptions about colour perception, since intricate empirical work is required to estimate model parameters (Kelber *et al*., 2017; Olsson *et al*., 2015; Vorobyev & Osorio, 1998). However, we contend that, on the contrary, some of these ‘simpler’ methods actually make very strong latent assumptions, which are not supported by the empirical evidence. This includes the assumption that all cones contribute equally to colour perception, that colour discrimination is unequivocal (i.e. the magnitude of colour-difference does not affect discriminability) and that colour differences follow an interval scale (as opposed to a ratio scale). Thus, we suggest that considering detectability relative to a threshold is essential for tests of discriminability. We emphasise, however, that this does not necessitate the use of the receptor-noise model specifically. Although we have focused on this popular approach here, particularly due to its utility for non-model organisms, a breadth of available modelling tools are capable of offering similar, and in some cases superior, insight (Kemp *et al*., 2015; Price & Fialko, 2017; Renoult *et al*., 2017). The hexagon model of Chittka (1992), for example, has been extensively tested and validated in honeybees, and may outperform the receptor-noise model when suitably parameterised (Garcia *et al*., 2017). It too offers a psychophysiologically-informed measure of perceptual distance, as well as discrimination thresholds, and so may be readily applied within our suggested framework. The two-step approach we propose can be easily and directly extended to these models.

Our simulations show that both the distance PERMANOVA and Cartesian MANOVA perform similarly well in statistically differentiating colours in perceptual space (Fig. 3). Studies have pointed out that distance-based methods perform poorly when the experimental design is unbalanced or when there is heteroscedasticity (though, among distance-based methods, PERMANOVA outperforms other approaches; Anderson & Walsh, 2013; Warton *et al*., 2012). It is important to note that these are often common features of, and applicable to, colour data (Endler & Mielke, 2005), and that these assumptions should be considered and verified. However, this might still be the most robust option for high-dimensional visual systems (e.g. Arikawa *et al*., 1987; Cronin & Marshall, 1989), by reducing data to a single metric of distance. Recently, Delhey *et al*. (2015) advocated a similar MANOVA approach, by applying a Principal Component Analysis (PCA) to the noise-corrected Cartesian coordinates prior to the test. However, if all the principal components are used in the MANOVA, results will be numerically identical to directly using the *XYZ* coordinates (which is preferable, since it is often tempting to discard PC axes of low variance, which could be problematic given that those axes may be involved in group differentiation). While we have focused on tests of differences in the location of colours in colour space, we recognise that other characteristics — such as differences in dispersion and correlation structure, and to identify the direction of variation among groups — might themselves be of biological interest, for which a PCA approach may be particularly useful.

The MANOVA approach can be extended to multivariate generalizations of generalized linear models by using the noise-corrected Cartesian coordinates as response variables (Hadfield, 2010). These models can also relax the assumptions of heteroscedasticity by estimating the variance-covariance of the responses (Had-field, 2010), and can be extended to include various error and model structures, such as hierarchical and phylogenetic models (Hadfield & Nakagawa, 2010). Still, these approaches will only test for the statistical separation in colour space, so estimating the magnitude of that separation is still necessary. The bootstrapped distance provides an easy to interpret measure of uncertainty to the mean distance estimate. Under a Bayesian approach, the mean distance bootstrap can be substituted by estimating credible intervals for the distance between perceptually-corrected Cartesian centroids from the posterior distribution, though this will be influenced by the priors adopted (Hadfield, 2010, see Electronic Supplementary Material for an example analysis).

Irrespective of the method used, it is essential to parametrize the underlying visual model appropriately (Garcia *et al*., 2017; Olsson *et al*., 2017). The Weber fraction and receptor densities chosen will strongly affect noise-corrected distances since they directly scale with the JND unit (Bitton *et al*., 2017). Further, even under adequate values of the Weber fraction it is important to realize that the unit JND usually reflects psychophysical performance under extremely controlled conditions (Kelber *et al*., 2003; Olsson *et al*., 2015), and that more conservative estimates of 2-4+ JND may be more appropriate for ecological and evolutionary questions (Osorio *et al*., 2004; Schaefer *et al*., 2007). Sensitivity analyses are also useful for exploring the robustness of conclusions against parameter variation, particularly in the case of non-model systems where such values are often assumed or drawn from related species (Bitton *et al*., 2017; Olsson *et al*., 2017). More broadly, we affirm recent (and ongoing) calls for pragmatism when drawing inferences from any such model (Marshall & Simmons, 2017; Olsson *et al*., 2017; Vasas *et al*., 2017). Colour spaces are valuable tools, but ultimately demand ongoing feedback from physiological and behavioural tests to improve our understanding of complex biological phenomena.

Our results show that insight into the biology of colour and its role in communication is best achieved by disentangling the implicit assumptions in questions of discriminability. By bringing these assumptions to light, our two-step approach offers a flexible procedure for examining the statistical presence and theoretical magnitude of differences between colour samples. We expect it will bring exciting new perspectives on the role of colour in intra- and interspecific interactions, and provide an efficient analytical framework for the study of colour in nature.

## Implementation and data accessibility

Analyses and simulations can be found in https://github.com/rmaia/msdichromatism/, and the described methods are fully implemented in the R package pavo as of version 1.3.1, available via CRAN. Key functions include bootcoldist which calculates the bootstrapped confidence intervals for mean distances, while jnd2xyz converts chromatic distances in JNDs to noise-corrected Cartesian coordinates. Multi-dimensional plotting options for noise-converted coordinates are also available. Lizard colour data from Whiting *et al*. 2015 are available at http://dx.doi.org/10.6084/m9.figshare.1452908.

## Acknowledgments

We would like to thank Ruchira Somaweera for leaf-nosed lizard photographs, and Dan Noble, John Endler and two anonymous reviewers for valuable discussion and comments on earlier drafts of the manuscript. This work was partially supported by a Junior Fellow award from the Simons Foundation to R.M., and an Australian Research Council grant (DP140140107) to T.E.W.

## Author contributions

RM and TEW conceived the ideas, designed methodology, analysed the data, and wrote the manuscript.

